# Spliceosomal miR-99b Regulates SPACA6-AS1 Pre-mRNA Levels and Promotes Malignant Phenotypes in Breast Cancer

**DOI:** 10.64898/2026.08.19.745696

**Authors:** Aya Muharram, Maram Arafat, Michal Linial, Ruth Sperling

## Abstract

MicroRNAs (miRNAs) are small non-coding RNAs that regulate gene expression primarily in the cytoplasm. However, emerging evidence highlights their additional roles in the nucleus. In particular, spliceosomal miRNAs have been implicated in novel regulatory functions, including the modulation of gene expression. Here, we investigate the nuclear role of spliceosomal miR-99b in breast cancer cells, focusing on its interaction with the long non-coding RNA (lncRNA) SPACA6-AS1. Using non-tumorigenic (MCF-10A) and breast cancer cell lines (MCF-7 and MDA-MB-231), we demonstrate that spliceosomal miR-99b expression increases with malignancy and correlates with elevated SPACA6-AS1 pre-mRNA levels. Notably, miR-99b exhibits full complementarity to the 5′ splice junction of SPACA6-AS1, suggesting a direct role in splicing regulation. Functional assays reveal that inhibition of miR-99b reduces SPACA6-AS1 pre-mRNA levels, whereas its overexpression enhances pre-mRNA accumulation, indicating that miR-99b promotes the formation or stabilization of the unspliced transcript. Furthermore, increased miR-99b expression is associated with altered ratios of SPACA6 isoforms, supporting a broader role in RNA-level regulation of gene expression. Phenotypically, miR-99b enhances breast cancer cell migration and is required for efficient invasion, particularly in highly aggressive cancerous cells. Our findings uncover a novel nuclear function of miR-99b in modulating lncRNA splicing and gene expression. This spliceosomal miR-99b–SPACA6-AS1 axis represents a previously unrecognized regulatory pathway that contributes to breast cancer progression and may provide a potential target for diagnostic and therapeutic strategies.

## 1. Introduction

MicroRNAs (miRNAs) are small non-coding RNAs, approximately 22 nucleotides in length, that play essential roles in regulating cellular signaling pathways. Canonically, miRNAs control gene expression post-transcriptionally by base-pairing with the 3′ untranslated region (3′-UTR) of target mRNAs in the cytoplasm, leading to translational repression or mRNA degradation [1–3]. Dysregulation of miRNA expression is associated with numerous chronic diseases and is fundamental to maintaining cellular identity and homeostasis. Changes in miRNA expression profiles are a hallmark of many cancer types. Based on their expression patterns in cancerous tissues compared to matched normal tissues, miRNAs can be broadly classified into two functional groups: (i) oncogenic miRNAs (oncomiRs), which promote tumor progression, and (ii) tumor-suppressive miRNAs, which inhibit cancer development [4–7].

The detection of miRNAs in the nucleus has expanded their functional landscape beyond cytoplasmic regulation [4,8,9]. Notably, both mature miRNAs and pre-miRNA-derived sequences have been identified within the supraspliceosome, suggesting additional nuclear roles [1,10–12]. Although these functions are not yet fully understood, accumulating evidence indicates that nuclear miRNAs participate in diverse processes, including the regulation of non-coding RNAs (ncRNAs), transcriptional silencing and activation, and modulation of transcriptional activity [13–16].

miRNAs have also emerged as key players in cancer biology, with growing interest in their use as diagnostic and prognostic biomarkers [17]. Breast cancer, one of the most commonly diagnosed malignancies and a leading cause of cancer-related mortality among women worldwide [18–22], has been extensively studied in this context. Numerous miRNAs have been implicated in breast cancer progression. However, their full mechanistic roles remain incompletely defined [5,7]. Representative oncogenic miRNAs include miR-10b, miR-21, miR-135a, miR-155, miR-221/222, miR-224, miR-373, and miR-520c [23–28], while tumor suppressors include the let-7 family, miR-7704, miR-30a, miR-31, miR-34a, miR-125 family, miR-200 family, miR-203, miR-205, miR-206, and miR-342 [10,23,29–32].

In our previous study [10], we analyzed spliceosomal miRNAs in breast cancer cell lines (MCF-7 and MDA-MB-231) and non-tumorigenic MCF-10A cells. We found that spliceosomal miRNA profiles are cell-type specific and differ from those reported in patient cohorts, suggesting distinct nuclear targets. We further demonstrated a negative regulatory interaction between spliceosomal miR-7704 and the oncogenic lncRNA HAGLR, supporting a tumor-suppressive role for nuclear miRNAs.

Recent analyses in neuronal and brain cancer models have similarly revealed novel nuclear targets of spliceosomal miRNAs, including miR-99b, which can enhance the levels of the lncRNA SPACA6-AS1 (Sperm acrosome associated 6-antisense 1) pre-mRNA through base pairing at its 5′ splice junction [12]. miR-99b is a conserved member of the miR-99 family (miR-99a/100/99b) that broadly functions as a context-dependent regulator of tumor biology, most commonly acting as a tumor suppressor through repression of key oncogenic pathways [33]. Mechanistically, miR-99b frequently targets components of the mTOR signaling axis, as well as genes involved in cell cycle progression, DNA repair, and epithelial–mesenchymal transition (EMT), thereby limiting proliferation, invasion, and therapy resistance [34–37]). In multiple solid tumors, including breast, prostate, lung, and hepatocellular carcinoma, miR-99b is typically downregulated and its restoration suppresses tumor growth and metastasis, consistent with inhibitory effects on pathways such as PI3K/AKT/mTOR signaling [34,36]. In contrast, in certain hematologic malignancies and specific tumor contexts, miR-99b can exhibit oncogenic or pro-survival roles, reflecting cell-type– specific target repertoires and regulatory networks [6,38]. Clinically, altered miR-99b expression highlights an emerging role as a biomarker and a potential therapeutic modulator in cancer [39]. Given the tissue-specific roles of miRNAs and their involvement in cancer-related pathways, understanding nuclear functions of miRNA is essential [40,41].

Here, we investigate the crosstalk between spliceosomal miR-99b and the lncRNA SPACA6-AS1 in breast cancer cells. These transcripts originate from the same genomic locus but are transcribed in opposite directions. We show that miR-99b directly increases SPACA6-AS1 pre-mRNA levels, likely through interaction with its splice junction. Functionally, miR-99b promotes breast cancer cell migration and invasion, while its inhibition suppresses these phenotypes. Additionally, miR-99b modulates the ratio among SPACA6 isoforms, linking nuclear miRNA activity to regulation of alternative transcript isoforms. Together, our findings identify a novel miRNA-lncRNA regulatory axis involving spliceosomal miR-99b and SPACA6-AS1, highlighting a previously unrecognized nuclear mechanism that contributes to breast cancer progression.

## 2. Results

### 2.1. Correlation between SF-miR-99b, SPACA6P-AS1 pre-mRNA, and breast cancer malignancy level

We previously identified differential expression of spliceosomal fraction miRNAs (SF-miRNAs) across breast epithelial cell lines spanning increasing malignant potential: MCF-10A (non-tumorigenic), MCF-7 and MDA-MB-231. Notably, their expression profiles differed from those described in the literature, raising the possibility that SF-miRNAs engage distinct nuclear targets and contribute to regulatory mechanisms separate from their established cytoplasmic activities [10]. Among these, miR-99b (within miR-99b/let-7e/miR-125a cluster) emerged as a compelling candidate (**Figure 1A**). miR-99b is encoded within intron 1 of a specific isoform of the SPACA6 gene, which, although primarily associated with the sperm acrosome, according to GTEx data indicate that it is also expressed at low levels in non-germline tissues and cell lines, including breast cancer cell lines. Importantly, the mature miR-99b-5p sequence is fully complementary to the 5′ splice site of intron 1 of SPACA6-AS1, a poorly characterized antisense (AS) long noncoding RNA (lncRNA) transcribed from the same locus in the opposite orientation (**Figure 1B**), and previously associated with adverse breast cancer prognosis [42].

**Figure 1.**
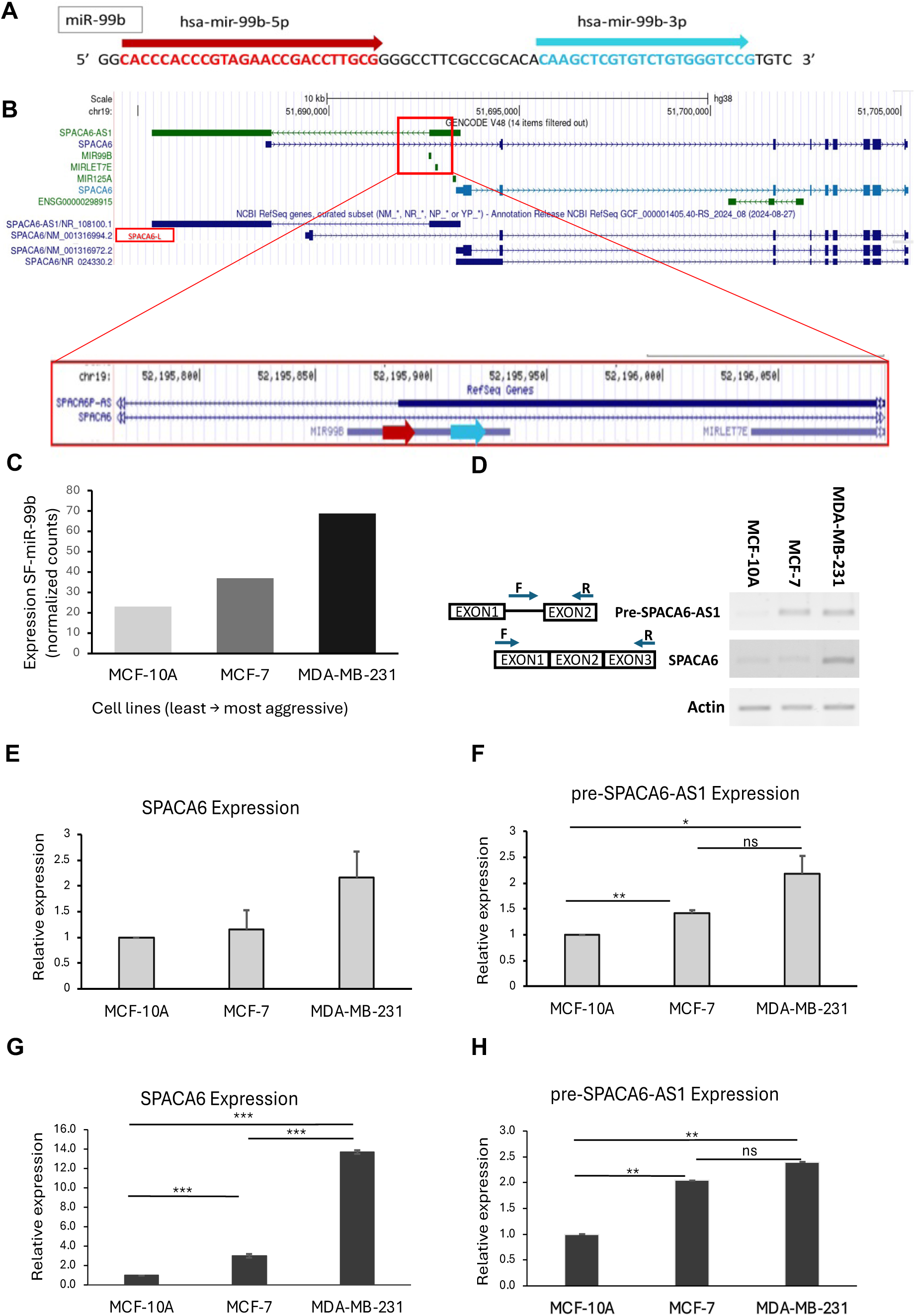
Direct correlation between the expression level of SF-miR-99b and SPACA6P-AS1 pre-mRNA in breast cancer cells malignancy. (A, B) The genomic location of miR-99b illustrates how it might affect the splicing of SPACA6-AS1. (A) The pre-miRNA sequence is depicted, with the 5p and 3p regions highlighted in red and blue, respectively. The graphical illustration centers around the isoforms of the SPACA6 gene. (B) Visualization of UCSC Genome browser (hg38) from Chr19 covering SPACA6-AS1 and the isoforms of SPACA6. The miR-99b cluster is indicated by the red frame and zoom-in around the miR-99b shows that miR-99b-5p can fully complement the 5′ splice junction of the intron of SPACA6-AS1, which is transcribed in the opposite direction of SPACA6, through base-pairing interaction. (C) Changes in the level of spliceosomal miR-99b in the breast cancer cell lines, as determined by RNA-Seq. (D) Breast cancer cell-lines (from MCF-10A, non-malignant, to malignant MCF-7 and metastatic MDA-MB-231) total RNA was analyzed by RT-PCR for the changes in expression of pre-SPACA6-AS1 and SPACA6. RT-PCR products were electrophoresed on a 2% agarose gel. Their identities, confirmed by sequencing, are given on the left; open boxes represent exons, lines represent introns, and arrow heads represent the PCR primers. actin, was used for normalization. (E) Quantification of RT-PCR results of total RNA SPACA6. (F) Quantification of RT-PCR results of total RNA SPACA6-AS1 pre-mRNA. (G, H) qRT-PCR analysis of the changes in expression of nuclear SPACA6 and SPACA6-AS1 pre-mRNA, respectively, in the three cell lines. The RT-PCR results of both total RNA and nuclear RNA reveal that the increase in the level of spliceosomal miR-99b is accompanied by significant increase in the level of SPACA6-AS1 pre-mRNA and increase in SPACA6 mRNA. Values represent the mean ± SEM of at least three independent experiments. Statistical significance was determined using unpaired two-tailed Student’s t-tests; not significant(ns), p < 0.05 (*), p < 0.01 (**).

RNA-seq analysis revealed a progressive increase in SF-miR-99b abundance with increasing malignancy (**Figure 1C**). Given its complementarity to the SPACA6-AS1 splice junction, we hypothesized coordinated regulation at the level of RNA processing. Consistent with this hypothesis, RT-PCR analysis of total RNA demonstrated increased levels of SPACA6-AS1 pre-mRNA in malignant cells compared with MCF-10A, with the highest levels observed in MDA-MB-231 (**Figures 1D, 1F**). A similar trend was observed for SPACA6 transcripts (**Figures 1D, 1E**). Analysis by qRT-PCR further confirmed progressive increase in nuclear SPACA6-AS1 pre-mRNA and SPACA6 RNA levels with malignancy (**Figures 1G, 1H**).

We also analyzed by RT-PCR the expression of SPACA6-AS1 (exon1, exon1) compared to SPACA6-AS1 pre-mRNA in nuclear RNA from the normal and breast cancer cell lines (**Figures 2A-C**). Consistent with the above findings, SPACA6-AS1 pre-mRNA levels increased with malignancy. In contrast, analysis of SPACA6-AS1 RNA (exon1, exon1), capturing both precursor and mature transcripts, revealed a different pattern, with maximal expression in MCF-7 and minimal levels in MDA-MB-231 (**Figures 2A, 2C**). Thus, the most aggressive cells MDA-MB-231 exhibit elevated miR-99b and SPACA6-AS1 pre-mRNA levels but reduced mature SPACA6-AS1 RNA, consistent with impaired splicing (**Figure 2D**), Indeed, the increased ratio between pre-mRNA/mature + pre-mRNA of SPACA6-AS1 in MDA-MB-231 relative to MCF-10A (**Figure 2D**) supports a model in which miR-99b negatively regulates SPACA6-AS1 splicing via direct interaction with its 5′ splice site. Collectively, these results indicate a direct association between the levels of SF-miR-99b and SPACA6-AS1 pre-mRNA, although contributions from additional members of the miR-99b cluster cannot be excluded.

**Figure 2.**
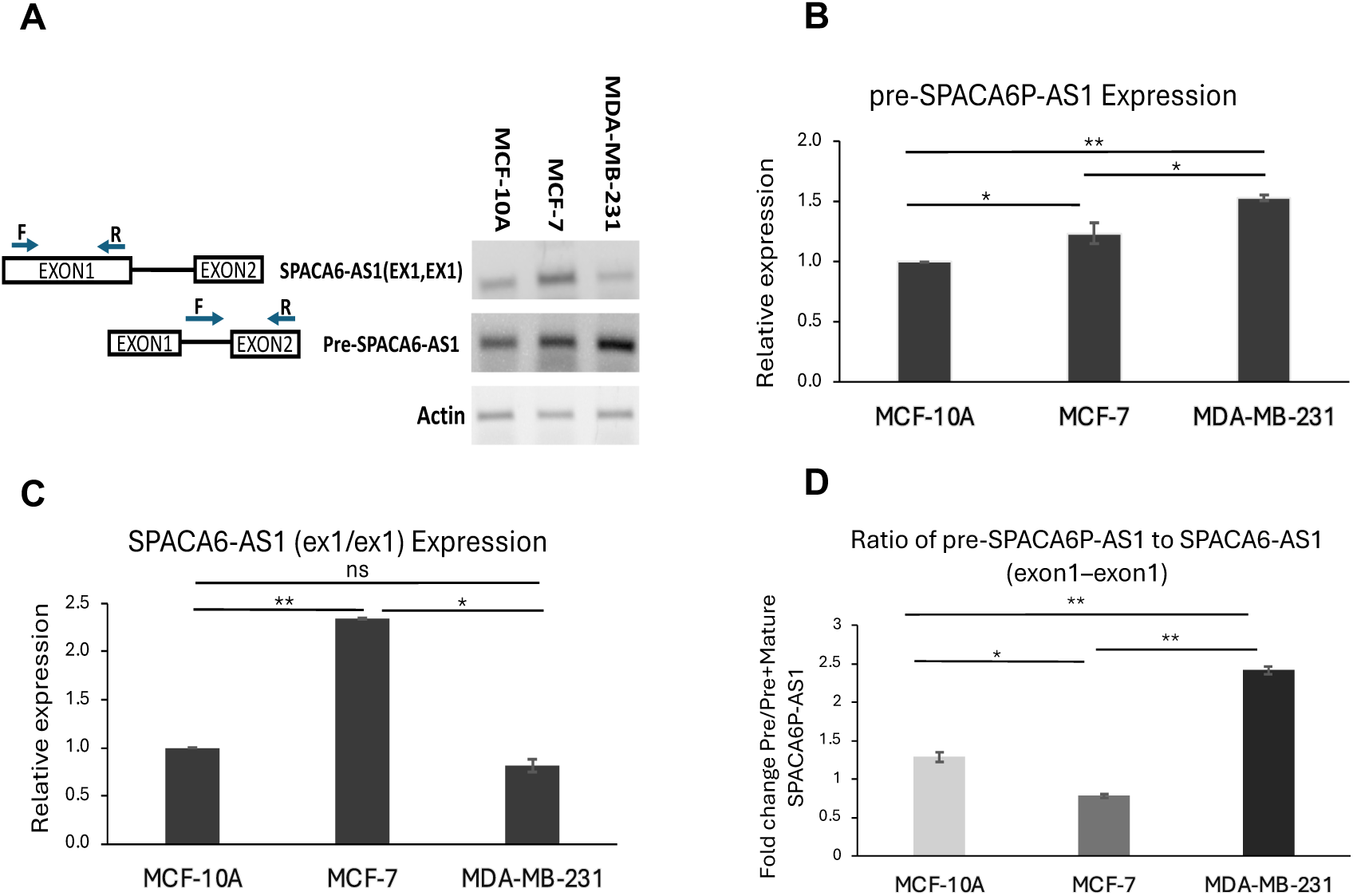
Changes in the expression of SPACA6-AS1 between normal and breast cancer cells. (A) Breast cancer cell-lines were analyzed by RT-PCR for the changes in expression of nuclear pre-SPACA6-AS1 and SPACA6-AS1(exon1, exon1) capturing both precursor and mature transcripts. RT-PCR products were electrophoresed on a 2% agarose gel. Their identities, confirmed by sequencing, are given on the left; open boxes represent exons, lines represent introns, and arrow heads represent the PCR primers. actin, was used for normalization. (B) RT-PCR analysis of RNA extracted from nuclear fractions of the normal and breast cancer cells for the pre-SPACA6-AS1. (C) RT-PCR analysis of RNA extracted from nuclear fractions of the normal and breast cancer cells for the total expression of SPACA6-AS1 (precursor and mature). (D) Relative ratios of un-spliced vs all SPACA6-AS1 splice isoforms, across the three cell lines, as determined in B and C, respectively. Values represent the mean ± SEM of at least three independent experiments. Statistical significance was determined using unpaired two-tailed Student’s t-tests; not significant(ns), p < 0.05 (*), p < 0.01 (**), p < 0.001 (***).

### 2.2. SPACA6-AS1 pre-mRNA expression is proportional to SF-miR-99b levels

To further investigate the relationship between miR-99b expression and SPACA6-AS1 pre-mRNA, breast cell lines were transfected with an anti-miR-99b inhibitor. Nuclear RNA was subsequently isolated, and the expression levels of miR-99b, SPACA6-AS1 pre-mRNA, and SPACA6 transcripts (**Figure 3A-C**, respectively) were quantified by qRT-PCR using TaqMan Fast Advanced Master Mix (Thermo Fisher Scientific) and gene-specific TaqMan assays. Cells transfected with a non-targeting anti-miR served as controls. Inhibition of miR-99b resulted in a consistent reduction in SPACA6-AS1 pre-mRNA levels across all three breast cell lines (**Figure 3B**). A concomitant decrease in SPACA6 expression was also observed (**Figure 3C**).

**Figure 3.**
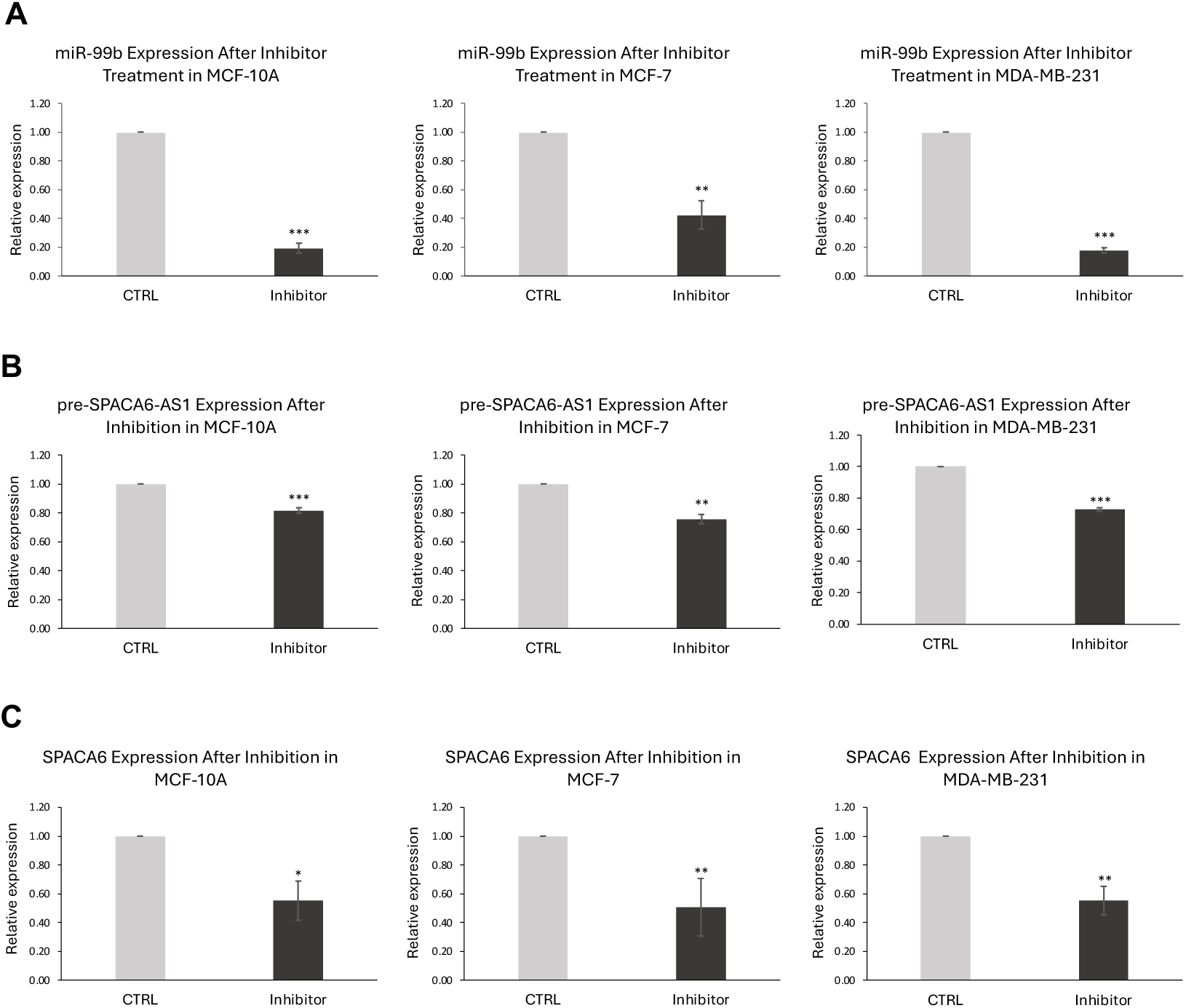
Inhibition of miR-99b downregulates the expression level of SPACA6-AS1 pre-mRNA. qPCR analysis of the effect of anti-miR-99b on the expression level of SPACA6-AS1 and SPACA6. (A) Transfection with anti-miR-99b downregulates the expression level of SF-miR-99b in normal and breast cancer cells. On the left are the control cells with anti-miR negative control (gray), and on the right are the results of cells transfected with miR-99b inhibitor (black). (B) Downregulation of the expression of SPACA6-AS1 pre-mRNA in breast cancer cells after inhibition of miR-99b (black), compared to the control (gray), (C) Downregulation of the expression of SPACA6 mRNA in breast cancer cells after inhibition of miR-99b (black), compared to the control (gray). Data represent mean ± SEM from at least three independent experiments. Statistical significance was determined using unpaired two-tailed Student’s t-tests; not significant(ns), p < 0.05 (*), p < 0.01 (**), p < 0.001 (***).

To further validate this relationship, miR-99b was overexpressed via transfection with a synthetic miR-99b mimic (**Figure 4A**) and nuclear RNA was isolated. As shown in **Figure 4B**, miR-99b overexpression led to a significant increase in SPACA6-AS1 pre-mRNA levels. Similarly, SPACA6 expression was elevated across all cell lines under these conditions (**Figure 4C**). These results demonstrate a direct and positive association between spliceosomal miR-99b levels and the expression of SPACA6-AS1 pre-mRNA, indicating that modulation of nuclear miR-99b is sufficient to influence SPACA6-AS1 transcriptional or post-transcriptional regulation.

**Figure 4.**
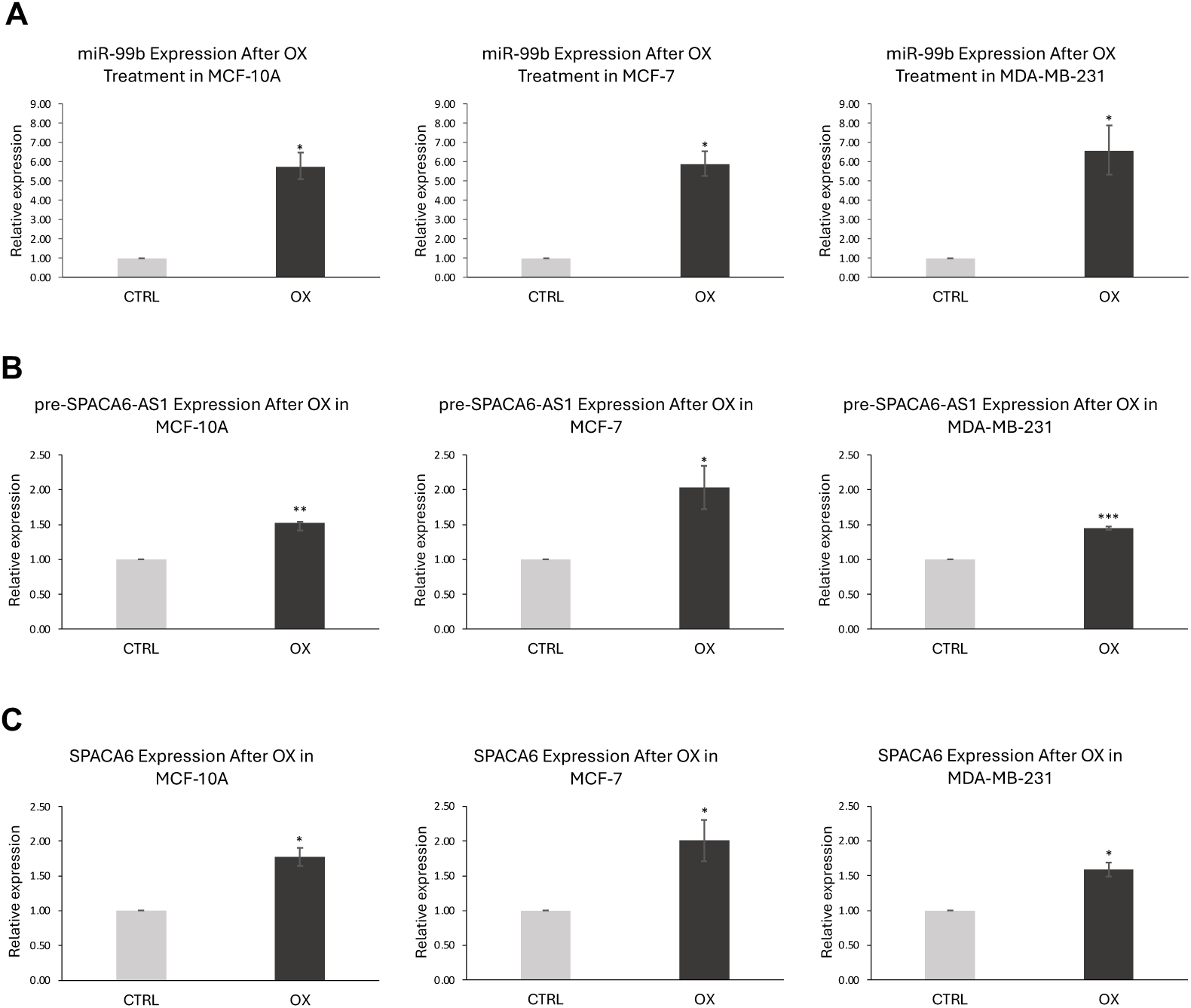
Overexpression of miR-99b is accompanied by increase in the expression level of SPACA6-AS1 pre-mRNA. qPCR analysis of the effect of overexpression of miR-99b on the expression level of SPACA6-AS1 and SPACA6. (A) Transfection with miR-99b mimic increases the expression level of SF-miR-99b in normal and breast cancer cells. On the left are the control cells miR mimic negative control (gray), and on the right are the results of cells transfected with miR-99b mimic (black). (B) Upregulation of the expression of SPACA6-AS1 pre-mRNA in breast cancer cells after overexpression of miR-99b (black), compared to the pre-miR negative control (gray), (C) Upregulation in the expression of SPACA6 mRNA in breast cancer cells after overexpression of miR-99b (black), compared to the control (gray). Data represent mean ± SEM from at least three independent experiments. Statistical significance was determined using unpaired two-tailed Student’s t-tests; not significant(ns), p < 0.05 (*), p < 0.01 (**), p < 0.001 (***).

### 2.3. Nuclear interaction network: miR-99b modulates SPACA6-AS1 pre-mRNA and SPACA6 isoform balance

SPACA6 is expressed as multiple transcript isoforms (**Figure 1A**), including NM_001316994.2 (hereafter termed SPACA6-L), which overlaps with a portion of an intron of the antisense lncRNA SPACA6-AS1 transcribed from the same genomic locus in the opposite orientation. Based on this genomic arrangement, we hypothesized that increased levels of SPACA6-AS1 pre-mRNA may influence SPACA6 isoform expression. To test this, we quantified SPACA6-L expression using qPCR primers spanning exons 1 and 3, and measured total SPACA6 (SPACA6-T) transcripts using primers targeting exons 2 and 3 (exons shared by all isoforms). In untreated breast epithelial cell lines, qPCR analysis revealed that both SPACA6-L and SPACA6-T expression increased in accordance with the degree of cells’ malignancy (**Figure 5A, B**). Notably, SPACA6-L constitutes approximately 10% of SPACA6-T transcripts in MCF-10A and increases to 19% in the aggressive MDA-MB-231 cell line (**Figure 5C**).

**Figure 5.**
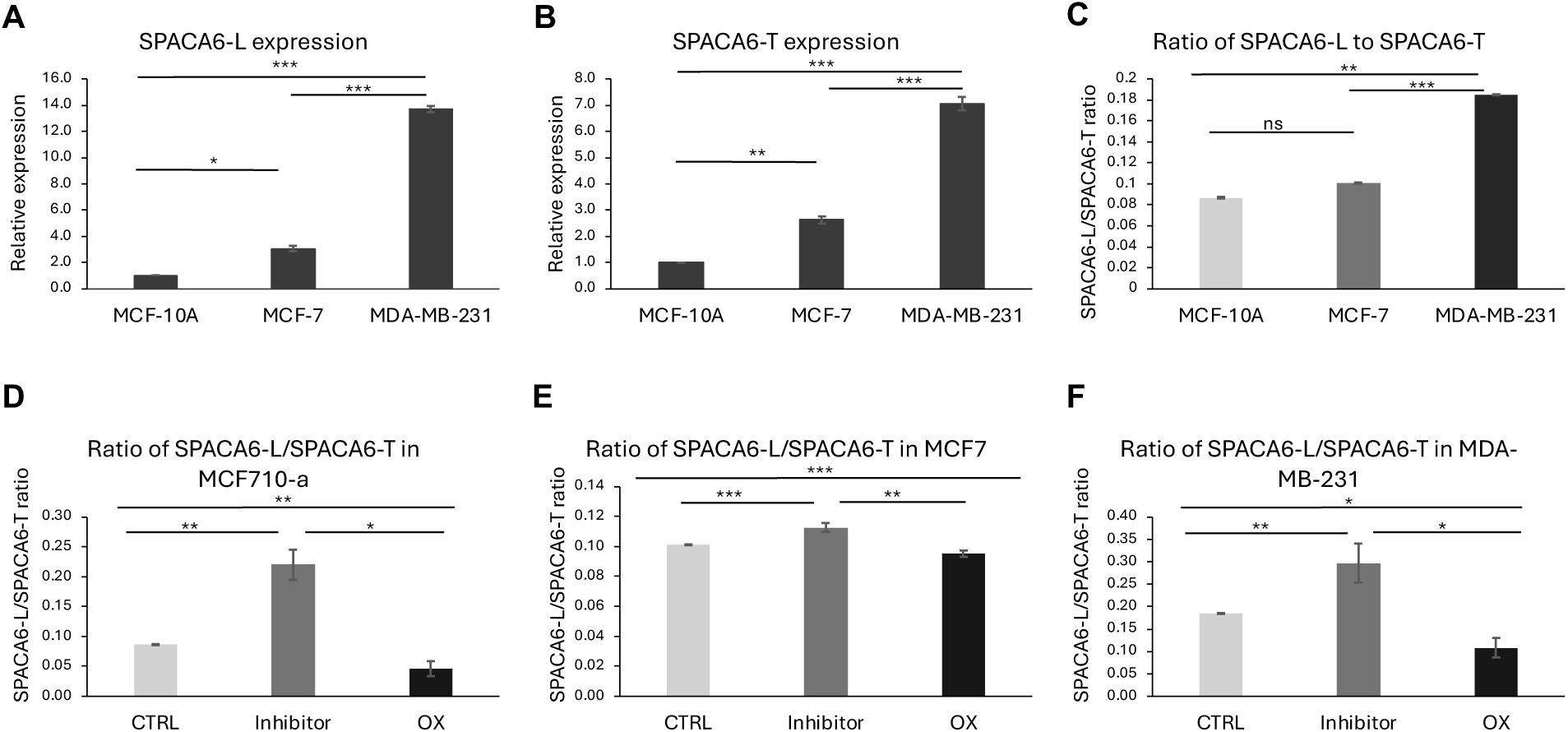
Changes in expression of SPACA6 isoforms in the breast cancer cell-lines – qPCR results. (A, B) qPCR analysis SPACA6-L and SPACA6-T in the untreated three breast cell lines, respectively. (C) qPCR analysis of the ratio between SPACA6-L/SPACA6-T in each of the untreated three breast cell lines. (D) qPCR analysis shows that miR-99b inhibition and overexpression affects the percentage of SPACA6-L in MCF-10A cells. (E) qPCR analysis shows that miR-99b inhibition and overexpression affects the percentage of SPACA6-L in MCF-7 cells. (F) qPCR analysis shows that miR-99b inhibition and overexpression affects the percentage of SPACA6-L in MDA-MB-231 cells. In untreated cells, SPACA6-L comprises 19% of total SPACA6. Upon inhibition of miR-99b (Inhibitor), the percentage of SPACA6-L increases to 30%, while overexpression of miR-99b decreased it to 11%. Data represent mean ± SEM from at least three independent experiments. Statistical significance was determined using unpaired two-tailed Student’s t-tests; not significant (ns), p < 0.05 (*), p < 0.01 (**), p < 0.001 (***).

To directly assess the role of miR-99b, we manipulated its levels and examined the resulting effects on SPACA6 isoform distribution. Inhibition of miR-99b, resulted in increased ratio of SPACA6-L with a similar trend in all cell lines, while overexpression of miR-99b resulted in decrease in SPACA6-L ratio, with a similar trend in all cell lines. Yet, the extent of changes in SPACA6-L percentage varied in the three breast cell lines, with minimal changes in the MCF-7 cell line, and increased changes in MCF-10A cell line (**Figure 5D-F**). The most significant changes occur in the aggressive malignant MDA-MB-231 cell line, where inhibition of miR-99b, associated with reduced SPACA6-AS1 pre-mRNA levels, led to an increase in the percentage of SPACA6-L transcripts from 19% to 30%, while, overexpression of miR-99b reduced the relative abundance of SPACA6-L to only ~11%. Together, these findings support a model in which miR-99b, through modulation of SPACA6-AS1 pre-mRNA, influences the balance of SPACA6 isoforms in the highly malignant cells.

### 2.4. Effect of miR-99b modulation on breast cancer cell migration

Cell migration and invasion are fundamental processes underlying metastatic progression. For tumor dissemination, cancer cells must acquire motility, invade through the extracellular matrix (ECM), intravasate into the circulation, and colonize distant sites. While migration assays assess intrinsic cell motility, invasion assays evaluate the capacity of cells to traverse ECM barriers, thereby modeling key aspects of metastasis.

To determine whether miR-99b regulates breast cancer cell motility, we performed wound-healing migration assays following miR-99b inhibition and overexpression in MDA-MB-231 and MCF-7 cells. As shown in **Figure 6A**, inhibition of miR-99b significantly reduced cell migration, as reflected by decreased relative wound density in both cell lines compared to controls, with a more pronounced effect observed in the highly aggressive MDA-MB-231 cells. Conversely, miR-99b overexpression (**Figure 6B**) enhanced cell migration, leading to increased relative wound density. This effect was statistically significant in MDA-MB-231 cells but not in MCF-7 cells, indicating a stronger functional impact in more aggressive cellular context. Collectively, these findings demonstrate that miR-99b promotes migratory capacity in breast cancer cell lines, with a more substantial effect in highly invasive phenotypes.

**Figure 6.**
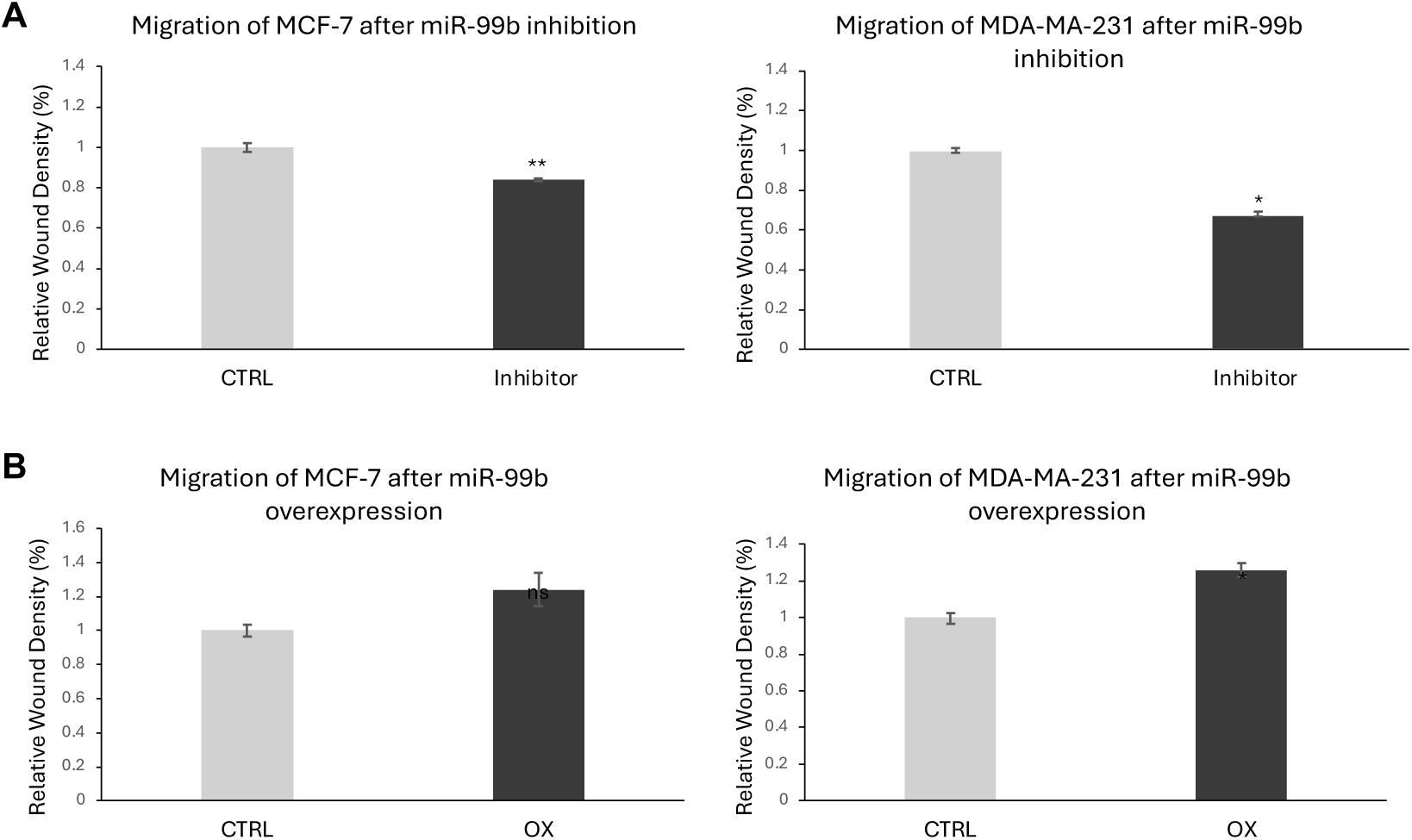
miR-99b increases the migration of breast cancer cells. (A) Migration of breast cancer cells (MDA-MB-231, MCF7) after 48 hours of treatment with miR-99b inhibitor. Wound-healing assays showed that downregulation of miR-99b reduced the migratory capacity of both cell lines, MDA-231 on the left, and MCF7 on the right, compared to control cells (CTRL) that were treated with non-silencing anti-miR negative control. (B) Migration of breast cancer cells (MDA-MB-231, MCF7) after 48h of treatment with miR-99b overexpression. Wound-healing assays showed that overexpression of miR-99b increased the migratory capacity of both cell lines, MDA-MB-231 on the left, and MCF7 on the right, compared to control cells (CTRL) that were treated with pre-miRNA negative control. Data represent mean ± SEM from at least three independent experiments. Statistical significance was determined using unpaired two-tailed Student’s t-tests: not significant (ns), p < 0.01 (*), p < 0.01 (**).

### 2.5. Effect of miR-99b modulation on breast cancer cell invasion

To assess the role of miR-99b in breast cancer cell chemotaxis and invasion through the ECM, we quantified invasion following miR-99b inhibition and overexpression, using relative wound density (%) as a readout. As shown in **Figure 7A**, inhibition of miR-99b significantly reduced invasive capacity in both MDA-MB-231 and MCF-7 cells compared to their respective controls, with a more pronounced effect observed in the highly aggressive MDA-MB-231 cell line. In contrast, overexpression of miR-99b did not enhance invasion in either cell line (**Figure 7B**), with only a modest (non-significant) increase detected in MCF-7 cells. These findings suggest that while miR-99b activity is required for maintaining invasive potential in breast cancer cells, its overexpression alone is insufficient to further augment invasion, indicating that additional factors or cooperative pathways are likely necessary to drive this cancer related phenotype.

**Figure 7.**
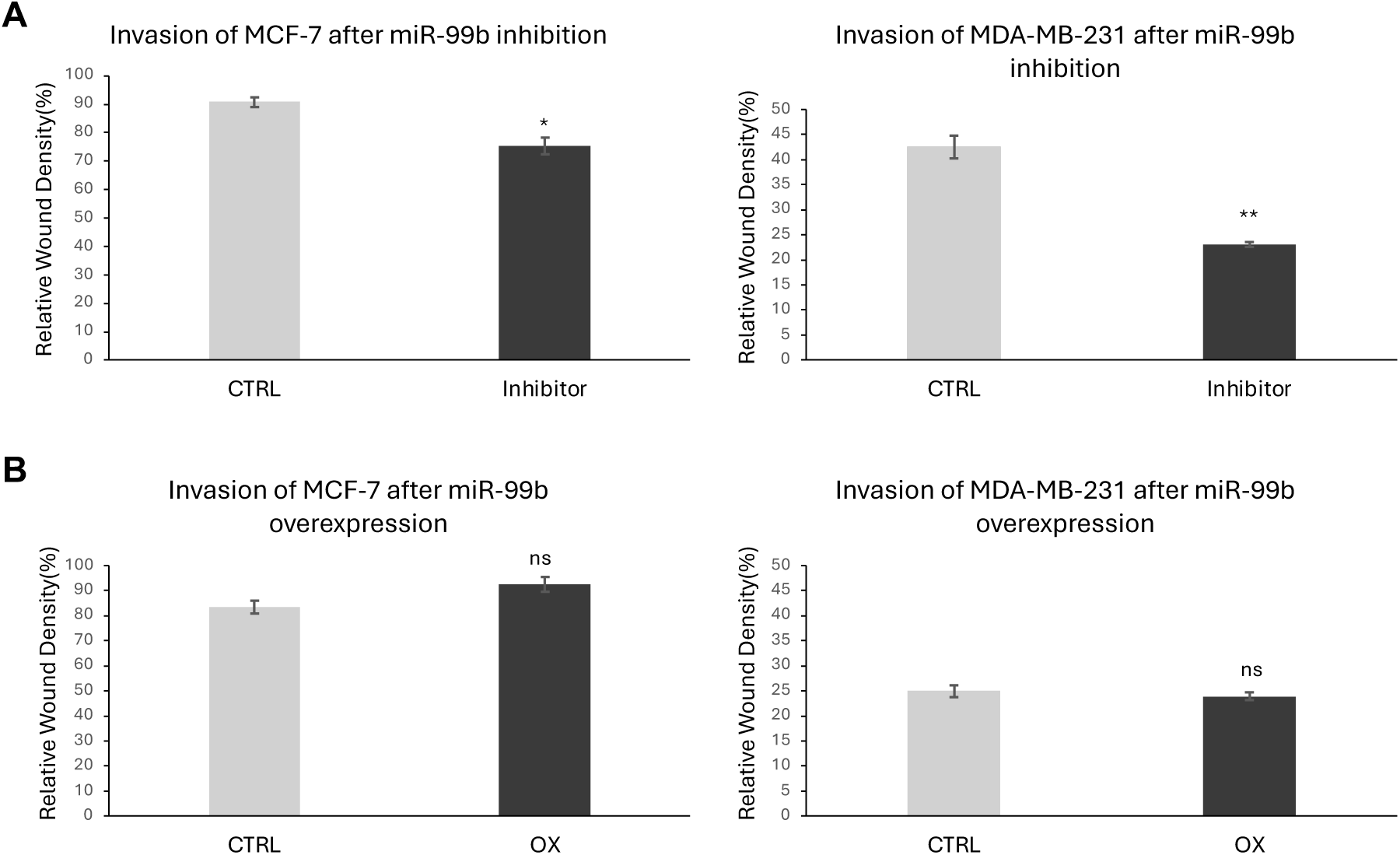
Effect of miR-99b inhibition and overexpression on the invasion of breast cancer cells. (A) Cell invasion of breast cancer cells (MDA-MB-231, MCF7) after 24h of treatment with miR-99b inhibitor. Wound-healing assays showed that downregulation of miR-99b reduced the invasion capacity of both cell lines, MDA-MB-231 on the left and MCF7 on the right, compared to control cells (CTRL) that were treated with non-silencing anti-miR negative control. (B) Cell invasion of breast cancer cells (MDA-MB-231, MCF7) after 24h of treatment with miR-99b overexpression. Wound-healing assays showed that overexpression of miR-99b doesn’t affect the invasion capacity of both cell lines, MDA-231 on the left and MCF7 on the right, compared to control cells (CTRL) that were treated with pre-miRNA negative control. Data represent mean ± SEM from three independent experiments. Statistical significance was determined using unpaired two-tailed Student’s t-tests: not significant (ns), p < 0.01 (*), p < 0.01 (**).

## 3. Discussion

miRNAs are well known for their canonical role in post-transcriptional gene regulation in the cytoplasm, where they inhibit translation or promote degradation of target mRNAs through base pairing with the 3′ untranslated region (3′-UTR) [1–3]. However, accumulating evidence has expanded this paradigm, demonstrating that miRNAs also function in the nucleus, where they regulate non-coding RNAs (ncRNAs), modulate transcription, and participate in transcriptional activation or silencing [4,8–16]. The detection of miRNAs and pre-miRNAs within spliceosomal complexes in HeLa, breast cancer, and neuronal cancer cells [1,10–12], including miRNAs not derived from introns, further supports functional roles for miRNAs in gene expression and RNA processing.

Genomic regions in which miRNAs are co-localized with protein-coding genes and antisense lncRNAs provide a structural basis for complex regulatory interactions. In such contexts, miRNAs may directly base pair with complementary sequences in antisense transcripts, forming regulatory networks that operate at the RNA level. For example, spliceosomal miR-7704 downregulates the oncogenic lncRNA HAGLR in HeLa and breast cancer cells [10,11]. Similarly, spliceosomal miR-99b has been shown to regulate SPACA6-AS1 pre-mRNA expression in neuronal systems [12] and in breast cancer cells, as demonstrated in this study (**Figures 1–4**). These findings emphasize the importance of exploring spliceosome-associated miRNAs as regulators of nuclear RNA networks and as potential therapeutic targets in cancer.

In this study, we focused on SF-miR-99b and its interaction with SPACA6-AS1 (also known as LINC01129), an antisense lncRNA located at chromosome 19q13.41. This locus is notable because SPACA6-AS1 overlaps antisense with the SPACA6 gene and with the miR-99b/let-7e/miR-125a cluster (**Figure 1**). Analysis of spliceosomal miRNA expression in breast cancer cell lines (MCF-7 and MDA-MB-231) compared to non-tumorigenic MCF-10A cells revealed that miR-99b levels increase with the degree of malignancy (**Figure 1C**). Importantly, miR-99b-5p is fully complementary to the 5′ splice junction of intron 1 of SPACA6-AS1 (NR_108100.1), spanning positions −6 to +16 relative to the splice site (**Figure 1B**). This complementarity suggests that spliceosomal miR-99b may interfere with the binding of U1 and U6 snRNP, required for the splicing process.

Consistent with this model, we observed a strong positive correlation between miR-99b levels and SPACA6-AS1 pre-mRNA accumulation across breast cancer cell lines. The most cancerous aggressive cells, MDA-MB-231, exhibited the highest levels of both miR-99b and SPACA6-AS1 pre-mRNA (**Figures 1C, 1D, 1F, 1H, 2A, 2B, 2D**), accompanied by reduced levels of mature+precursor of SPACA6-AS1 transcripts (**Figure 2C**), indicative of impaired splicing. This is further supported by the increased ratio of SPACA6-AS1 pre-mRNA/SAPACA6-AS1 (pre-mRNA+mRNA) (**Figure 2D**). Functional modulation of miR-99b reinforced this relationship. Namely, inhibition of miR-99b reduced SPACA6-AS1 pre-mRNA levels (**Figure 3B**), whereas overexpression increased its accumulation (**Figure 4B**). Together, these findings demonstrate that spliceosomal miR-99b directly regulates SPACA6-AS1 RNA processing, likely by inhibiting splicing and promoting accumulation of the un-spliced transcript.

Functionally, miR-99b contributes to breast cancer cell migration and invasion. Inhibition of miR-99b significantly reduced both migration and invasion in MDA-MB-231 and MCF-7 cells (**Figures 6A, 7A**), whereas overexpression enhanced migration (**Figure 6B**) but did not significantly increase invasion (**Figure 7B**). These results suggest that miR-99b is required for maintaining invasive capacity but is not sufficient on its own to further enhance invasion, indicating that additional regulatory factors are involved. Notably, these effects are more pronounced in the highly aggressive MDA-MB-231 cells, highlighting the context-dependent role of miR-99b.

We further show that modulation of miR-99b expression, which affects the expression of SPACA6-AS1 pre-mRNA, can also have an effect on SPACA6 isoforms. Modulation of miR-99b expression affects the ratio between SPACA6 isoforms, particularly in the highly malignant MDA-MB-231 cells (**Figure 5**). In these cells, inhibition of miR-99b increased the proportion of SPACA6-L from 19% to 30%, whereas overexpression reduced it to 11% (**Figure 5F**). Given that SPACA6-AS1 pre-mRNA partially overlaps with SPACA6-L in an antisense orientation, these findings suggest that increased SPACA6-AS1 pre-mRNA levels may influence SPACA6-L levels, and in the more malignant MDA-MB-231 cells also influences isoform balance, potentially through RNA–RNA interactions.

The observation that the percentage of SPACA6-L in the MDA-MB-231 cell-line is higher than in the MCF-10A and MCF-7 cells, is likely explained by the contribution of additional factors, such as the expression level of SF-miR-125a and SF-Let-7e.

Although SPACA6 is primarily known as a sperm-associated protein involved in sperm–egg fusion, its role in breast cancer cells remains unclear. Similarly, the biological function of SPACA6-AS1 is still poorly characterized. According to GTEx data, its expression is highest in testis but is also detectable at lower levels in other tissues, including breast cancer cells. Notably, SPACA6-AS1 has been identified as a risk-associated lncRNA inversely correlated with breast cancer survival [42], suggesting a potential oncogenic role. Our findings extend this observation by implicating the pre-mRNA form of SPACA6-AS1 as an active component of a nuclear regulatory network involving miR-99b and SPACA6 transcripts. Additional complexity may arise from other members of the miR-99b cluster, including miR-125a and let-7e. For example, in hepatocellular carcinoma, SPACA6-AS1 participates in a competing endogenous RNA (ceRNA) network with miR-125a and its targets (Lin28b, MMP11, SIRT7, ZBTB7A), where reciprocal regulation between SPACA6-AS1 and miR-125a influences oncogenic pathways [43]. miR-125a itself is a tumor suppressor that is downregulated in cancer [23,44] and negatively regulates the RNA-binding protein HuR, which is associated with cell proliferation, migration, and poor prognosis in breast cancer [44]. These observations suggest that the miR-99b/SPACA6-AS1 axis may be functionally linked to broader nucleocytoplasmic regulatory networks, although inspecting this possibility remains beyond the scope of the current study.

The identification of multiple spliceosomal miRNAs that overlap with antisense lncRNAs across ovarian [1,11], breast [1,10], and neuronal cancer systems [12] supports the existence of nuclear RNA regulatory networks mediated by RNA-RNA interactions. Similar to the regulation of HAGLR by miR-7704 [10], our findings demonstrate that miR-99b modulates SPACA6-AS1 splicing and expression (**Figures 1–4**) and additionally influences SPACA6 isoform balance (**Figure 5**). In conclusion, we describe a novel nuclear regulatory axis involving spliceosomal miR-99b, SPACA6-AS1, and SPACA6 isoforms in breast cancer cells. Through direct base pairing with a splice junction (**Figure 1B**), miR-99b likely modulates lncRNA splicing (**Figures 1–4**), alters transcript isoform distribution (**Figure 5**), and promotes cancer cell migration and invasion (**Figures 6, 7**). These findings expand the functional repertoire of miRNAs and suggest that nuclear miRNA-RNA interactions represent a previously underappreciated layer of gene regulation with potential therapeutic relevance.

## 4. Materials and Methods

### 4.1. Cells

RNA was isolated from the following cell lines: MCF-10A, non-tumorigenic human mammary epithelial cell line; MCF-7 mammary epithelial adenocarcinoma cell line and MDA-MB-231 aggressive metastatic tumorigenic human mammary epithelial cell line.

#### 4.1.1. Cellular modulation of miR-99b levels (inhibition and overexpression)

After 24h of seeding the cells (MCF-10A, MCF-7, and MDA-MB-231) in six-well plates, the medium was changed, and two tubes were prepared for effective transfection. The first tube contains 150 µL of Opti-MEM medium with 9 µL of Lipofectamine RNAiMAX Reagent (cat# 13778075, ThermoFisher), and the second tube contains 150 µL of Opti-MEM medium with 3 µL (10 µM) of miRNA. After mixing the tubes and incubating at room temperature for 5 minutes, 250 µL of this solution was added to each well.

For downregulation of hsa-miR-99b, cells were transfected with 10 μM of hsa-miR-99b Anti-miR inhibitor (cat #AM17000, Ambion). For control experiments cells transfected with the Anti-miR negative control (cat#AM17010, Ambion). Cells were collected after 24h for downstream analysis. For the setting of overexpression of hsa-miR-99b, 24h before nuclear RNA isolation was performed (described below) the cell lines were transfected with 10 μM of the hsa-miR-99b Pre-miR Precursor (cat #AM17100, Ambion). For control, we used transfection with Pre-miR Negative control (cat# AM17110, Ambion).

### 4.2. RNA extraction

#### 4.2.1. Total RNA

Each type of cell was seeded in six-well plates that were washed with PBSx1, followed by the addition of 300 µL of Trypsin 0.25%–EDTA (L0931-500, Biowest) per well, transferred to a tube, and centrifuged for 3 min at 300g. Total RNA was extracted using the RNeasy Mini Kit (Qiagen), according to the manufacturer’s standard protocol.

#### 4.2.2. Nuclear RNA

Nuclear RNA isolation was performed as previously described [45]. Briefly, each type of cells was seeded in six-well plates, that were washed with PBSx1 followed by the addition of 175 µL of cold RLN buffer (50 mM Tris pH 8, 140 mM NaCl, 1.5 mM MgCl_2_ and 0.5% NP40) per well. The cells were then scraped and moved to an Eppendorf tube on ice for 5 min. Centrifugation for 2 min at 300g, at 4°C. The supernatant that contains the cells cytoplasm was transferred to a new tube, and the pellet (nuclei) was further processed. RNA was then extracted from nuclei fraction with miRNeasy mini kit (Qiagen), following the manufacturer’s instructions.

### 4.3. PCR analysis

RT-PCR analysis. RT-PCR was performed on total RNA extracted from the cells (MCF-10A, MCF-7, and MDA-MB-231) from the nuclear supernatants of the cells. The following sets of primers for SPACA6 were used: Forward (Fw, exon 1) 5′-CGTCGGAGCGGCAGAACTT-3′ and Reverse (Rev) 5′-TCATTTTCTCCGCAGCATC-3′ (exon 3) and run in PCR Tm 62°C for 35 cycles. The primers for pre-SPACA6-AS1 are: Fw (intron 1) 5′-GGGCTCAAAGGTGAATCAGA-3′ and Rev 5′-AGGGCCTTAGTGGAGGTCAT-3′ (exon 2) and run in PCR Tm 58 °C for 35 cycles. The primers for SPACA6-AS1(exon 1) are: Fw (exon 1) 5′-GCCCCCACCAGCTTTAGTATCT-3′ and Rev (exon 1) 5′-GTCTGCGGCTGGCTCTGT-3′ and run in PCR Tm 60 °C for 35 cycles. Sanger sequencing verified the identities of all PCR products. Each experiment was repeated at least 3 times. The relative abundance was quantified in view of the intensity of the β-actin. The β-actin Fw and Rev primers are: 5′-CTGGAACGGTGAAGGTGACA-3′ and 5′-AAGGGACTTCCTGTAACAATGCA-3′, respectively.

### 4.4. Quantitative PCR (qPCR)

#### 4.4.1. TaqMan microRNA Assay

For RT of miR-99b, the TaqMan MicroRNA Revers Transcription Kit (AB-43666596, ThermoFisher) was used according to the manufacturer’s instructions, using RT hsa-miR-99b primers, and MultiScribeTM Reverse Transcriptase. For PCR we used has-miR-99b Primers (TaqMan® MicroRNA Assays INV, SM/PC hsa-miR-99b:000436, AB-4427975) and MultiScribe™ Reverse Transcriptase (ThermoFisher).

#### 4.4.2. RT of mRNA

For the RT of mRNA, we performed RT of nuclear RNA using the High-Capacity cDNA Reverse Transcription Kit (AB-4368814, ThermoFisher) according to the manufacturer’s instructions, using RT Random Primers, and MultiScribeTM Reverse Transcriptase.

#### 4.4.3. Quantitative PCR reaction

mRNA and miR-99b levels were measured using the TaqMan Fast Advanced Mix (AB-4444557, ThermoFisher) and the following TaqMan Assays primers: TaqMan Advance MiRNA Assay hsa-mir-99b (AB-4427975, ThermoFisher); TaqMan Gene Expression Assays INV/SM: ACTB: Hs99999903_m1 (AB-4331182, ThermoFisher); Custom TaqMan Gene Expression Assays, SM/EA, SPACA6-AS1 intron-exon (AB-4331348, ThermoFisher). Primer Fw: 5′-GCCCCCACCAGCTTTAGTATCT-3′, Rev: 5′-GTCTGCGGCTGGCTCTGT-3′, FAM Probe: 5′-GGAGCTCACAGTCTGA-3′, SPACA6 total (exon 2-exon 3) (AB-4331348, ThermoFisher). Primer Fw: 5′-GAGAGAAGCCACCTGCATGAC-3′, Rev: 5′-CATTTTCTCCGCAGCATCAG-3′, FAM Probe: 5′-TCACCCAGATGACCCAT-3′. TaqMan Gene Expression for SPACA6 (NM_001316994.2 exon 1-exon 3 middle transcript-L): Custom TaqMan Copy Number Assays, SM/PC: ID APH6HV3 (AB-4400294, ThermoFisher). Assays were performed according to the manufacturer’s instructions.

A QuantStudio 3 Real-Time PCR System was used to amplify the sample for 40 cycles at 60°C annealing temperature. Analysis was performed using the 2^ΔΔCT method. At least three biological replications were used for each experiment.

### 4.5. Functional assays

#### 4.5.1. Migration assay

For preparing the migration assay plates, a thin layer of ECM GEL (E1270, Sigma) was applied to each well of a 96-well ImageLock plate (Essen BioScience, Cat# 4379). First, an aliquot of ECM was thawed and then diluted in 6 mL of complete media to a final concentration of 100 µg/mL. Next, 50 µL of the diluted ECM GEL was dispensed into each well of the 96-well ImageLock plate, ensuring the ECM GEL evenly coated each well. After coating, the plates were incubated overnight in a humidified incubator at 37°C with 5% CO₂ to allow the ECM GEL to solidify. Followed by aspiration of the ECM GEL from each well, breast cancer cells (MCF-7, and MDA-MB-231) were seeded into the assay plate and incubated overnight.

The confluent cell monolayers were wounded using the Incucyte 96-Well Woundmaker Tool. Followed by two washes with cell culture media at room temperature, media containing treatments for downregulation or overexpression of hsa-miR-99b at a final concentration of 10 μM were added to each well, and the plate was placed in the IncuCyte instrument. The IncuCyte software, repeat scanning every 2 hours over 48 hours using the Scratch Wound Wide Mode, when the first scan beginning immediately. The collected data from the scans was analyzed by the IncuCyte assay software to examine the effects of the treatments on cell migration.

#### 4.5.2. Invasion assay

Plate preparation with the ECM Gel and cell seeding was performed as above (Cell migration assay). Subsequently, the confluent cell monolayer was wounded using the Incucyte 96-Well Woundmaker Tool. After washing twice with cell culture media at room temperature, 100 µL of culture media were added to each well and the plate was placed in a pre-chilled CoolBox 96F to equilibrate for 5 min. Then, the media was removed, and 50 µL of 1 mg/mL ECM GEL solution and culture media per well were added to the assay plate and incubated for 30 min on the pre-warmed CoolSink 96F in a 37°C, 5% CO₂ environment. Subsequently, media containing downregulation or overexpression treatments of hsa-miR-99b at a final concentration of 10 μM were added to each well, and the plate was placed in the IncuCyte. In the IncuCyte software, repeat scanning every 2h over 24h using the Scratch Wound Wide Mode, with the first scan beginning immediately. The collected data from the scans was analyzed by the IncuCyte assay software to examine the effects of the treatments on cell invasion.

## Author Contributions

R.S. M.A. and M.L. conceived and designed the study. M.A. supervised the experiments; A.M. performed the experiments; A.M. M.A. and R.S.analysed the data; R.S. M.A and M.L. wrote the manuscript. All authors discussed the findings, have read the manuscript and agreed to the final version of the manuscript.

## Acknowledgments

We thank Aviva Petcho for excellent technical assistance, and Keren Zohar for support in the initial phase of this study. We thank the Center for Genomic Technologies at the Hebrew University and the Interdepartmental Equipment unit, Hadassah Ein Karem, at the Hebrew University. We gratefully acknowledge Prof. Maayan Salton at The Hebrew University of Jerusalem, Faculty of Medicine (Ein Karem Campus) for providing access to her laboratory equipment, including the biological safety cabinet, which was essential for this work.

## Abbreviations

AS: antisense
EMT: epithelial–mesenchymal transition
Fw: Forward primer
lncRNA: long non-coding RNA
miRNA: MicroRNA
qPCR: quantitative polymerase chain reaction
Rev: Reverse primer
SF: spliceosomal fraction

